# RAS mutation-specific signaling dynamics in response to paralog- and state- selective RAS inhibitors

**DOI:** 10.1101/2025.02.14.638317

**Authors:** Beau Baars, Ana Orive-Ramos, Ziyue Kou, Bijaya Gaire, Mathieu Desaunay, Christos Adamopoulos, Stuart A. Aaronson, Shaomeng Wang, Evripidis Gavathiotis, Poulikos I. Poulikakos

## Abstract

A high therapeutic index (TI), balancing potent oncogenic signaling inhibition in tumor cells with minimal effects on normal cells, is critical for effective cancer therapies. Recent advances have introduced diverse RAS-targeting inhibitors, including mutant-specific inhibitors (e.g., KRAS(G12C) and KRAS(G12D)), as well as paralog- and state-selective inhibitors. Non-mutant-specific RAS inhibition can be accomplished by 1) panRAS-GEF(OFF) inhibitors which inactivate RAS indirectly by inhibiting SHP2 or SOS1, thereby blocking the nucleotide exchange step of RAS activation, 2) direct KRAS(OFF)-selective inhibitors sparing NRAS and HRAS, and 3) panRAS(ON) inhibitors that directly target active RAS, by occluding binding of its effector RAF. However, the signaling inhibition index (SII) - the differential inhibition of oncogenic signaling between RAS-mutant (RAS(MUT)) and normal cells - remains poorly defined for these approaches.

In this study, we evaluated the SII of state- and paralog-selective RAS inhibitors across diverse RAS-mutant (RAS(MUT)) and RAS-wild-type (RAS(WT)) models. PanRAS-GEF(OFF) inhibitors exhibited neutral or negative SII, with comparable or reduced MAPK suppression in KRAS(G12X) cells relative to RAS(WT) cells. KRAS(G13D) models showed low sensitivity (negative SII) to panRAS-GEF(OFF) inhibitors, particularly in the context of NF1 loss. Combination treatments with SHP2 and MEK inhibitors resulted in low SII, as pathway suppression was similar in RAS(MUT) and RAS(WT) cells. Furthermore, RAS(Q61X) models were resistant to combined SHP2 inhibitor+MEK inhibitor due to dual mechanisms: MEK inhibitor-induced NRAS(Q61X) reactivation and RAS(MUT)-induced SHP2 conformations impairing inhibitor binding.

Overall, panRAS-GEF(OFF) inhibitors exhibited the lowest SII. PanKRAS(OFF) inhibitors demonstrated a higher SII, while panRAS(ON) inhibitors displayed broader activity but relatively narrow SII. We observed that tumors that were sensitive to RAS(MUT)-specific inhibitors, were also sensitive to the state-selective RAS inhibitors (OFF, or ON). In fact, all RAS inhibitors (mutant-specific and state- or paralog-selective) were active in the same portion of RAS(MUT) models, while the majority of RAS(MUT) cell lines were insensitive to all of them. These findings reveal significant SII variability among RAS-targeted inhibitors, depending on the specific RAS driver mutation and cell context and underscore the importance of incorporating SII considerations into the design and clinical application of RAS-targeted therapies to improve therapeutic outcomes.

**Main points:** *PanRAS-GEF(OFF) inhibitors have limited SII and effectiveness:* The Signaling Inhibition Index (SII) – i.e. the differential inhibition of oncogenic signaling between tumor and normal cells - was neutral or negative for panRAS-GEF(OFF) inhibitors, with comparable or reduced MAPK suppression in KRAS(G12X) mutant versus RAS(WT) cells. KRAS(G13D) models showed reduced sensitivity, particularly with NF1 loss. SHP2+MEK inhibitor combinations also had low SII, with RAS(Q61X) models demonstrating resistance due to NRAS(Q61X) reactivation and impaired SHP2 inhibitor binding.

*PanKRAS(OFF) selective inhibitors have higher SII than panRAS-GEF(OFF) inhibitors:* panKRAS(OFF)-selective inhibitors have a higher SII compared to panRAS-GEF(OFF) inhibitors, offering better tumor-versus-normal cell selectivity.

*PanRAS(ON) inhibitors have broad but modest SII:* While panRAS(ON) inhibitors displayed a broader activity profile, their ability to selectively inhibit mutant RAS signaling over normal cells remained relatively narrow (low SII).

*Most KRAS-mutant tumors will be insensitive to any single RAS-targeted inhibitor:* State- and paralog-selective inhibitors have enhanced activity in the same RAS-MUT cancer models that are also sensitive to RAS-MUT-specific inhibitors, suggesting that most KRAS-MUT tumors will not respond uniformly to any one RAS-targeting inhibitor.

*SII varies across RAS inhibitors, necessitating tailored therapeutic strategies:* The effectiveness of paralog- and state-selective inhibitors depends on specific RAS mutations and cell context, highlighting the need to integrate SII considerations into the development and clinical application of RAS-targeted therapies.

## Introduction

The RAS family (KRAS, NRAS, and HRAS) of small GTPases regulates MAPK signaling and other downstream effectors by cycling between GTP-bound (active, ON) and GDP-bound (inactive, OFF) states^1,2^. Under normal conditions, RAS signaling is tightly regulated and essential for cell proliferation, differentiation, and survival. This regulation depends on the coordinated activity of GEFs, such as SOS, which promote GTP loading, and GTPase-Activating Proteins (GAPs), such as NF1, which facilitate GTP hydrolysis. Activated RAS recruits RAF to the membrane for activation of the kinase cascade RAF/MEK/ERK. Activated ERK promotes a transcriptional program that drives cell cycle progression. RAS proteins are frequently mutated in human cancer^1-3^, with most common mutations occurring in positions G12, G13 and Q61^1,3^. These mutations shift the steady-state equilibrium in cells towards the RAS(ON) state to varying degrees, driving uncontrolled cell proliferation and tumorigenesis^1,3^.

Despite being discovered over four decades ago, RAS(MUT) has historically been considered “undruggable” due to the lack of well-defined binding pockets for small molecules^4^. However, pioneering work by the Shokat group^5^, led to the development of the first direct inhibitors targeting KRAS(G12C)^6-8^, exploiting a druggable pocket in its inactive (GDP-bound or OFF) state. This breakthrough has spurred the development of numerous small-molecule inhibitors targeting KRAS(G12C)^9-12^, and more recently, KRAS(G12D)^13^.

Contrary to the long-held belief that RAS(MUT) are “locked” in the active (GTP-bound) state, recent studies^14-17^, including our own^18^, have demonstrated that RAS(G12X) mutants exist in a dynamic equilibrium between the GTP-bound (ON) and GDP-bound (OFF) states. This discovery has expanded the therapeutic potential of small molecules targeting the inactive RAS conformation, either indirectly, by targeting SHP2^14,16-19^ or SOS1^20^ and consequently the nucleotide exchange step of RAS activation (**Fig. 1A**), (panRAS-GEF(OFF) inhibitors-panRAS-GEF(OFF)i), or directly, in a paralog-selective manner, e.g, inhibitors that target both KRAS(WT) and KRAS(MUT), while sparing NRAS and HRAS^21^ (panKRAS(OFF) inhibitors-panKRAS(OFF)i). Recent advancements have also led to the development of direct, panRAS(ON) inhibitors (panRAS(ON)i), also known as RAS(ON) multi-selective inhibitors, (e.g. RMC-7977, ERAS-0015, GFH547), which bind Cyclophilin A (CYPA) to form a high-affinity complex that sterically occludes RAS–effector interactions^22^. This approach enables the targeting of active RAS, a previously inaccessible conformation, and has shown activity across a broad spectrum of RAS(MUT)^22,23^. One such compound, RMC-6236, is currently in Phase I clinical trials (NCT05379985).

**Figure 1.**
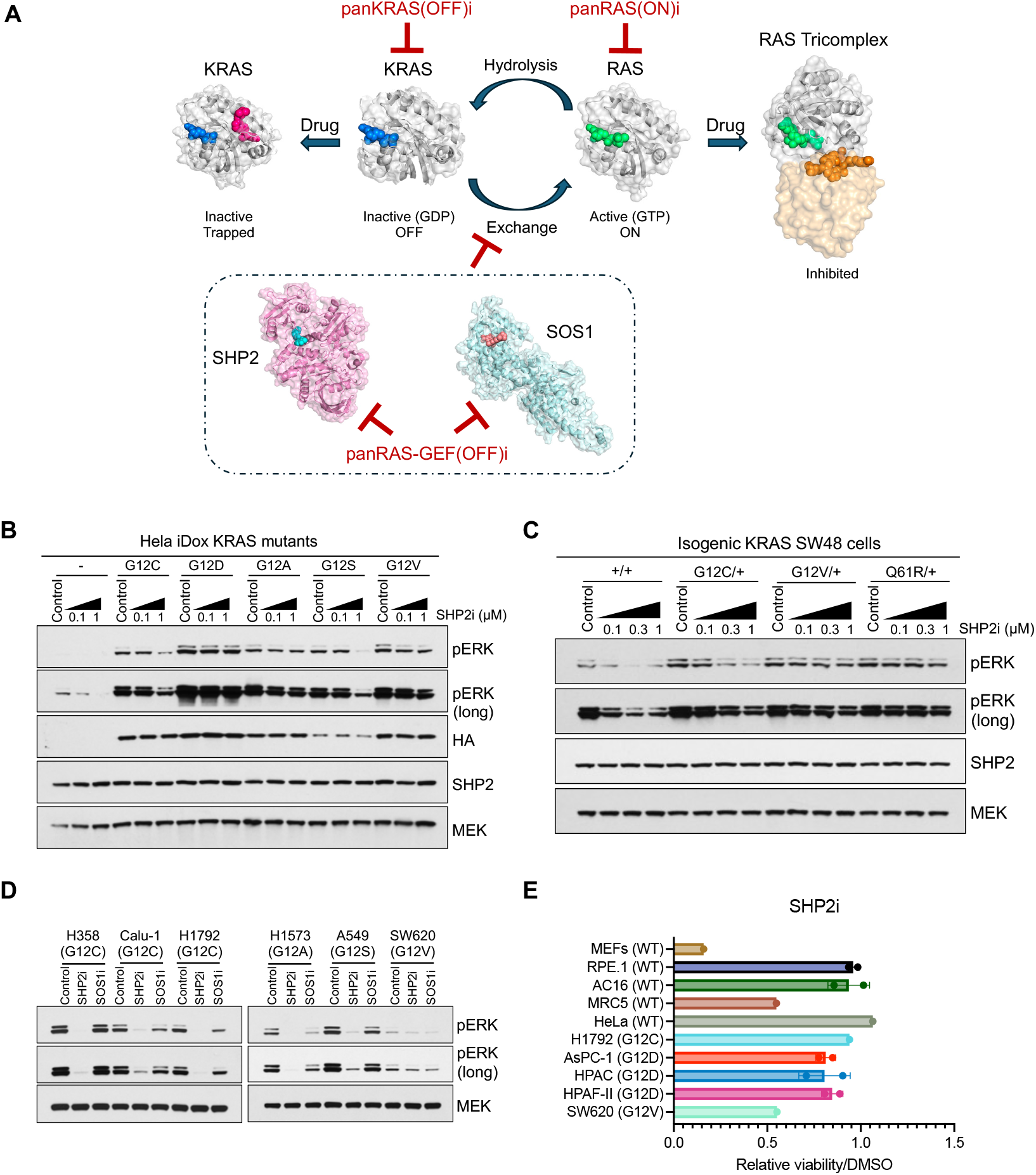
KRAS(G12X)-expressing cells exhibit differential sensitivity to panRAS-GEF(OFF)i. **(A)** RAS cycles between inactive/OFF (GDP (blue)-bound) state and active/ON (GTP (green)-bound) state under the control of SHP2 and Guanine Exchange Factors (GEFs), such as SOS1. The inactive/OFF state can be targeted indirectly by panRAS-GEF(OFF)i which block SHP2 and SOS1 from promoting RAS activation, or inhibitors that directly inhibit KRAS in the OFF state (panKRAS(OFF)i). The active/ON state of RAS can be targeted directly by panRAS(ON)i that forms a RAS tricomplex and prevents downstream MAPK pathway activation. Each panel is colored to highlight nucleotides/inhibitors. From left to right: KRAS bound to GDP (blue) and BI-2865 (pink) (Protein Databank (PDB): 8AZV), KRAS bound to GDP (blue) (PDB: 4OBE), KRAS bound to GTPγS, a non-hydrolysable nucleotide (green) (PDB: 5VQ6), KRAS bound to GppNHp, a non-hydrolysable nucleotide, along with RMC-7977 (orange) and cyclophilin A (orange) (PDB: 8TBK), SHP2 (pink) bound to TNO155 (light blue) (PDB: 7JVM), SOS1 (light blue) bound to BI-3406 (pink) (PDB: 6SCM). **(B)** Parental and Doxycycline-inducible KRAS variant HeLa cells were treated for 24 hours with increasing concentrations of doxycycline (50-600ng/mL), followed by treatment for 2 hours with SHP2i TNO155 (0.1 or 1 µM). SHP2 and MAPK pathway activation was monitored by immunoblotting. **(C)** Parental and isogenic SW48 cells expressing different KRAS(G12X) mutant were treated for 2 hours with SHP2i TNO155 (0.1, 0.3 and 1 µM). MAPK pathway activation was monitored by immunoblotting. **(D)** A panel of KRAS(G12X) cell lines were treated with either SHP2i TNO155 (5 µM) or SOS1i BI-3406 (10 µM) for 2 hours. MAPK pathway activation was assessed by monitoring pERK levels via immunoblotting. **(E)** The indicated cell lines expressing either KRAS(WT) or different KRAS(G12X) mutants were treated with SHP2i TNO155 (2 µM) for 6 days. The inhibitory effect on cell viability was monitored and presented as mean ± SEM from two independent experiments (n=2).

Despite these advancements, achieving a high Therapeutic Index (TI) remains a significant challenge in RAS/MAPK-targeted therapies. The Signaling Inhibition Index (SII), defined as the ratio of signaling inhibition in tumor versus normal cells, is a critical determinant of TI. While RAS(MUT)-specific inhibitors offer a theoretically unlimited SII, the SII of paralog- and state-selective RAS inhibitors remains poorly understood. In this study, we aimed to determine and compare the SII of state- and paralog-selective RAS inhibitors by evaluating signaling dynamics and tumor responses in RAS(MUT) and RAS(WT) models.

### KRAS(G12X)-expressing cells exhibit differential sensitivity to panRAS-GEF(OFF)i

To assess the sensitivity of different KRAS(G12X) mutations to panRAS-GEF(OFF)i, we utilized isogenic HeLa cells engineered with doxycycline (dox)-inducible expression of KRAS(G12C), KRAS(G12D), KRAS(G12A), KRAS(G12S), and KRAS(G12V) (**Fig. 1B**). SHP2 inhibitor (SHP2i) (chemical compounds used in this study are listed in **Fig. S1**) was highly potent against KRAS(G12C), KRAS(G12A) and KRAS(G12S), consistent with previous reports^14-16^ including ours^18^. However, SHP2i exhibited reduced activity in KRAS(G12D)- and KRAS(G12V)-expressing cells. Notably, SHP2i led to greater MAPK inhibition in RAS(WT)-expressing cells compared to KRAS(MUT) cells. We next tested SHP2i in RAS(WT) SW48 cells engineered to express KRAS(MUT) from the endogenous promoter using CRISPR knock-in (**Fig. 1C**). SHP2i was less potent in suppressing MAPK signaling in KRAS(G12V)-compared to KRAS(G12C)-expressing cells. MAPK signaling in KRAS(Q61R)-expressing cells was completely resistant to SHP2i, consistent with previous reports by us and others^16,18^. Similar to our experiments in HeLa cells, SHP2i potently suppressed MAPK in parental, RAS(WT) SW48 cells.

To further evaluate panRAS-GEF(OFF)i activity on KRAS(G12X) at the endogenous level, we tested treatment with SHP2i or SOS1i in a panel of KRAS(G12X) cell lines. The SOS1i used was consistently less potent than the SHP2i, while both showed a suppressive effect on MAPK in G12C/A/S KRAS(MUT) cells. In the KRAS(G12V)-expressing SW620 cells, basal MAPK levels were lower and minimally affected by either SHP2i or SOS1i (**Fig. 1D**).

Finally, we assessed the effect of SHP2i on cell viability of various KRAS(G12X) cells, along with RAS(WT) cell lines, modeling normal cells. SHP2i treatment did not universally suppress cell proliferation more potently in KRAS(G12X) over RAS(WT) cells. Two RAS(WT) cell lines showed substantial growth inhibition upon SHP2i treatment, whereas several KRAS(G12X) cells were only marginally affected (**Fig. 1E**).

Together, these findings indicate that SHP2i, when used as monotherapy, has a low SII, as its inhibitory effects are not sufficiently selective for KRAS(G12X) over RAS(WT) cells. This may explain the limited clinical efficacy observed in clinical trials assessing SHP2i for the treatment of RAS(G12X) cancers.

### NF1 Loss Confers Resistance to panRAS-GEF(OFF)i in KRAS(G13D) cells

Similar to KRAS(G12X) mutants, KRAS(G13D) mutants exhibit low but detectable intrinsic GTPase activity^24^, suggesting a dependence on upstream RTK signaling to maintain their active state. However, previous studies have reported conflicting findings regarding the sensitivity of KRAS(G13D) cells to SHP2i. While we previously demonstrated that SHP2i is ineffective in KRAS(G13D) models^18^, Mainardi et al. reported SHP2 dependence of KRAS(G13D)-mutant cells^15^, and Hofmann et al. showed sensitivity of a KRAS(G13D) line (DLD-1) to SOS1i^20^. SHP099^19^, the SHP2i used in our study^18^ is several-fold less potent than current clinical SHP2i. To confirm that the two KRAS(G13D) models we used in our study are also insensitive to next generation SHP2i and SOS1i, we treated the two KRAS(G13D) models, HCT-15 and HCT-116 previously used by us with the second generation SHP2i TNO155 or the SOS1i BI-3406. In both models, MAPK inhibition was minimal by either inhibitor (**Fig. S2**), consistent with our report^18^.

The KRAS(G13D) models tested by the various groups differ in their NF1 status: DLD-1 is NF1-proficient, whereas HCT-15 and HCT-116 used by us lack NF1 expression. This suggested that KRAS(G13D) cells require NF1 for SHP2 and SOS1 dependency. In support of this idea, either SHP2i or SOS1i treatment resulted in a modest suppression of MAPK signaling in NF1-proficient DLD-1 cells, but had no effect in NF1-deficient HCT-116 cells (**Fig. 2A**). To directly assess whether NF1 loss drives insensitivity to SHP2i in the KRAS(G13D) context, we knocked down NF1 using siRNA in DLD-1 (KRAS(G13D)) cells. NF1 downregulation resulted in complete resistance of MAPK inhibition by SHP2i (**Fig. 2B**), consistent with NF1 expression being required for sensitivity to SHP2i. Thus, KRAS(G13D) signaling is altered by NF1 expression, and NF1 loss diminishes sensitivity to panRAS-GEF(OFF)i. NF1 downregulation also promoted SHP2i insensitivity in a RAS(WT) model (**Fig. 2B**), suggesting that NF1 inactivation may be a general mechanism promoting insensitivity to SHP2i, in certain contexts.

**Figure 2.**
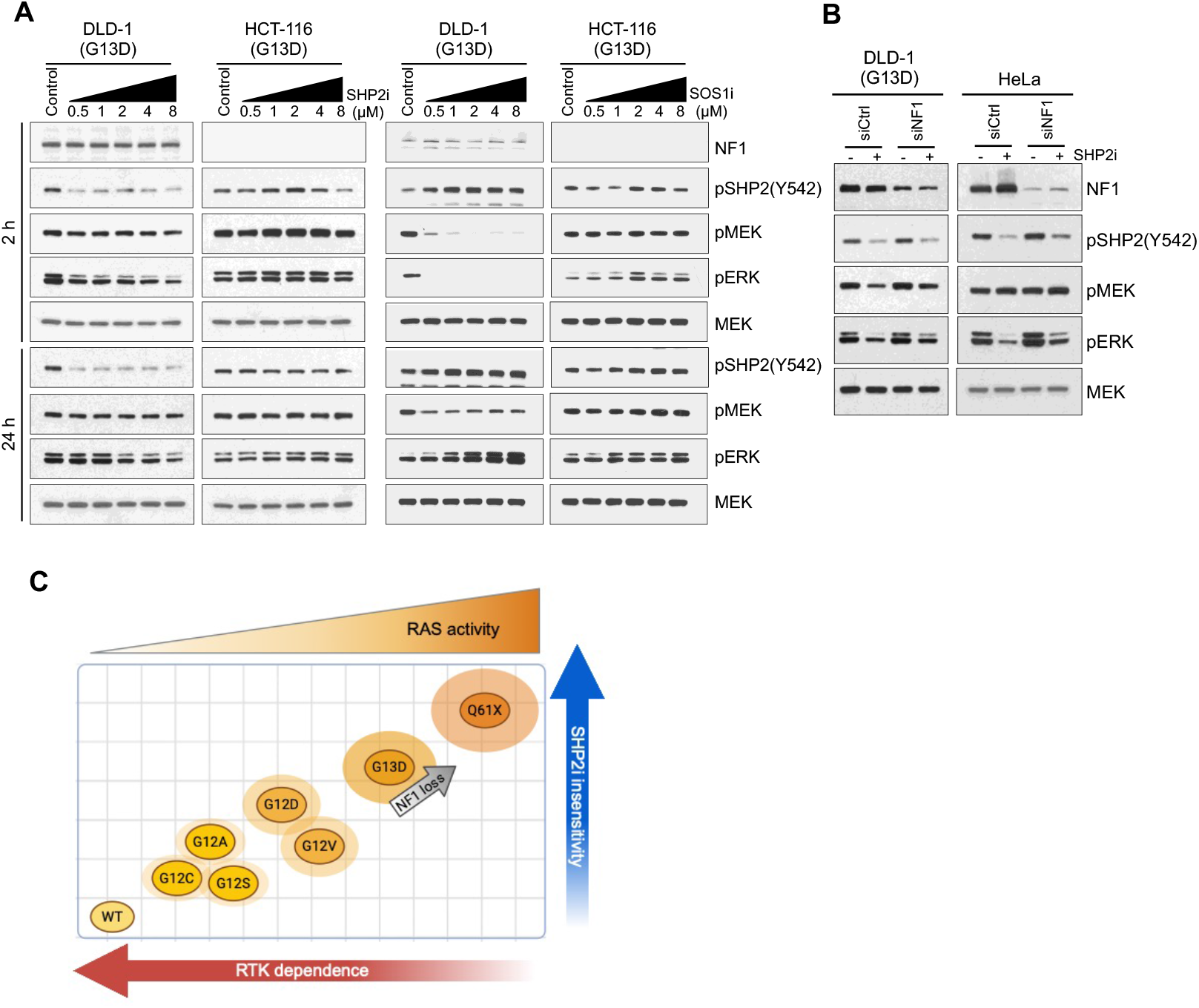
NF1 Loss Confers Resistance to panRAS-GEF(OFF)i in KRAS(G13D)-expressing cells. **(A)** KRAS (G13D)-mutant cell lines DLD-1 (NF1-proficient) and HCT-116 (NF1-deficient) were treated for 2 or 24 hours with increasing concentration (0.5, 1, 2, 4, 8 µM) of SHP2i TNO155 or SOS1i MRTX-0902. SHP2 and MAPK pathway activation were monitored by immunoblotting, respectively indicated by pSHP2^Y542^ and pMEK/pERK. **(B)** NF1 expression was transiently knocked-down in DLD-1 and HeLa, followed by treatment with SHP2i TNO155 (8 µM in DLD-1 and 4 µM in HeLa) for 2 hours. SHP2 and MAPK pathway activation were monitored by immunoblotting, respectively indicated by pSHP2^Y542^ and pMEK/pERK. **(C)** Schematic model depicting the ranking of KRAS(WT) and KRAS(MUT) variants based on their sensitivity to SHP2 inhibition, which correlates with the contribution of receptor tyrosine kinase (RTK)-mediated MAPK pathway activation. This figure was created in BioRender. Baars, B. (2025) https://BioRender.com/y27u304.

Together our findings show that RAS(MUT) vary in their sensitivity to panRAS-GEF(OFF)i. RAS(MUT) with increased dependence on upstream RTK/SHP2/SOS signaling have the highest sensitivity to SHP2i (WT, KRASG12C/S/A). KRAS(G12V/D) and NF1-proficient KRAS(G13D) have modest sensitivity to SHP2i, while NF1-deficient KRAS(G13D) and RAS(Q61R/K) cells, exhibit complete insensitivity to SHP2i (**Fig. 2C**).

### Combined SHP2i+MEKi suppresses MAPK activity in both normal and RAS(G12X) cells, diminishing tumor selectivity

MEK inhibitors (MEKi) have been assessed as a potential treatment for RAS(MUT) tumors, however adaptive resistance through negative feedback and RAS/MAPK reactivation limits their efficacy^25-27^. Thus, combining MEKi with SHP2i has been shown by us^18^ and others^14,15,28-30^ to enhance MAPK pathway suppression and mitigate resistance in various RAS(MUT) models.

We observed that SHP2i alone effectively inhibited MAPK activity in KRAS(G12C/A/S) cells, and to a lesser extent, in KRAS(G12V/D) models (**Fig. 1B, 1D**). To assess whether the SHP2i+MEKi combination could provide sustained MAPK suppression, we assessed pERK levels following short-term (2 hours) and long-term (24 hours) treatment. While SHP2i alone transiently reduced ERK activity in H1792 (KRAS(G12C)) cells, combined treatment of SHP2i plus MEKi resulted in more durable MAPK suppression (**Fig. 3A**). Similar results were observed when MEKi was combined with SOS1i (**Fig. 3A**). In KRAS(G12D) cells that were less sensitive to SHP2i monotherapy, MAPK was also potently inhibited upon combined SHP2i+MEKi treatment (**Fig. 3B**). To evaluate the tumor selectivity of this approach, we tested the SHP2i+MEKi combination in isogenic HeLa cells with doxycycline (dox)-inducible expression of RAS(WT) or various KRAS(G12X) mutants. SHP2i alone inhibited pERK more potently in RAS(WT) cells compared to KRAS(G12X) cells. Further, combined SHP2i+MEKi resulted in comparable MAPK suppression in both RAS(WT) and KRAS(G12X) cells, indicating lack of selectivity for the KRAS(G12X) over the RAS(WT) context (**Fig. 3C**).

**Figure 3.**
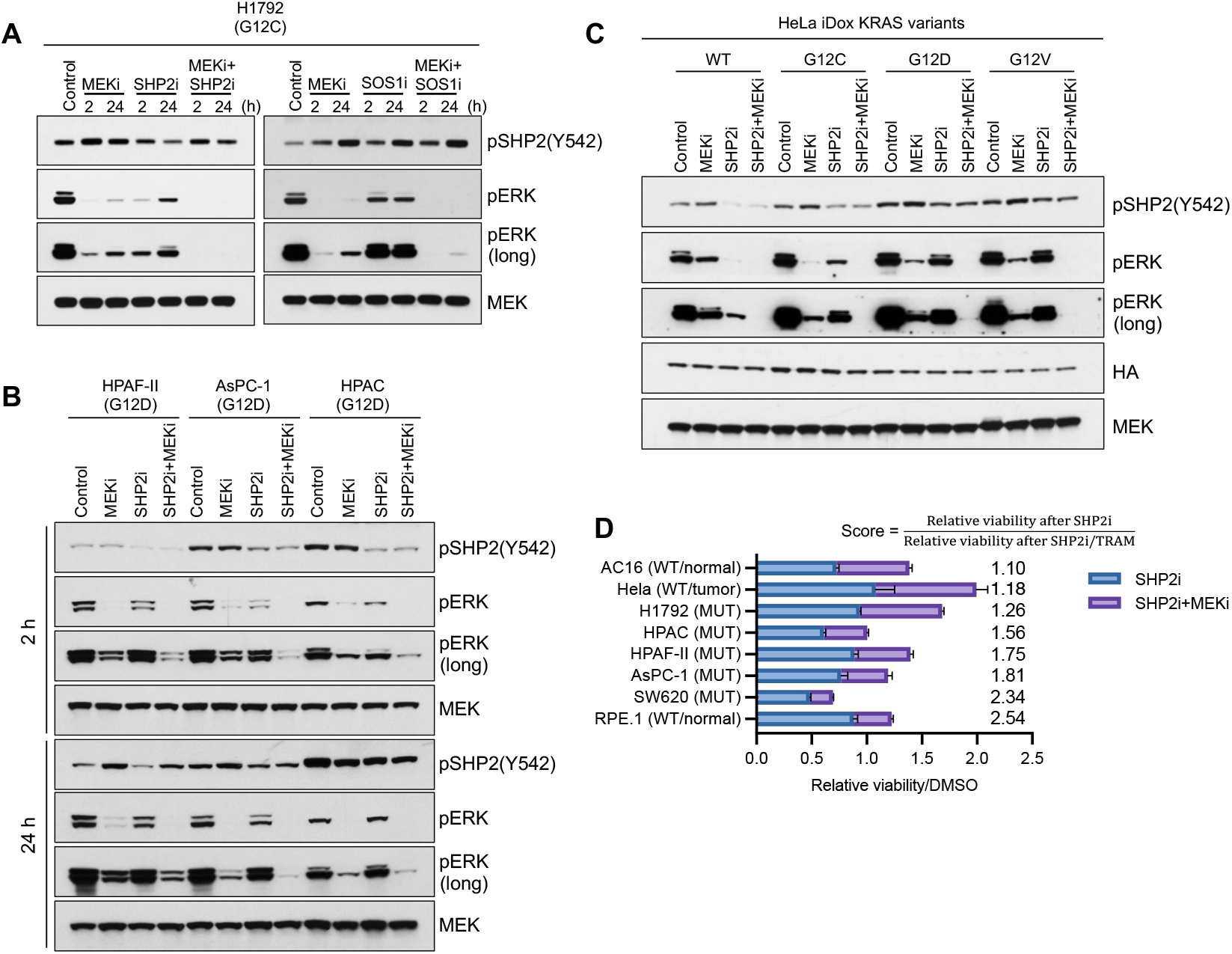
SHP2i+MEKi combination suppresses MAPK activity in both normal and KRAS(G12X)-mutant cells, diminishing tumor selectivity. (**A)** The indicated KRAS(G12X) cells was treated with 5nM MEK inhibitor trametinib, 5 μM SHP2i TNO155, 10 µM SOS1i BI-3406, or the combinations MEKi+SHP2i or MEKi+SOS1i. SHP2 and MAPK pathway activation were monitored by immunoblotting, respectively indicated by pSHP2^Y542^ and pERK. (**B**) The indicated KRAS(G12D) cells were treated with 5 nM MEK inhibitor trametinib, 2 μM SHP2i TNO155, or the combination MEKi+SHP2i. SHP2 and MAPK pathway activation were monitored by immunoblotting, respectively indicated by pSHP2^Y542^ and pERK. **(C)** Doxycycline-inducible KRAS variant HeLa cells were treated with increasing concentrations of doxycycline (50-600ng/mL) for 24 hours, followed by treatment with MEKi trametinib (5 nM), SHP2i TNO155 (2 µM), or the combination MEKi+SHP2i. SHP2 and MAPK pathway activation were monitored by immunoblotting, respectively indicated by pSHP2^Y542^ and pERK. **(D)** A panel of cell lines expressing either KRAS(WT) or different KRAS(MUT) was treated for 6 days with SHP2i TNO155 (5 µM) or SHP2i+ MEKi trametinib (3 nM). Cell viability inhibition was assessed and presented as a score assigned to each cell line, representing the ratio between the relative viability after SHP2i treatment and the relative viability after combination. Data are represented as mean values ± SEM, n=3 experimental replicates.

Finally, we assessed the effect on cell viability of SHP2i and SHP2i+MEKi in a panel of normal and tumor cell lines expressing RAS(WT) or KRAS(G12X), all of which were insensitive to SHP2i alone. To quantitatively compare the effects of SHP2i monotherapy versus SHP2i+MEKi treatment, we assigned viability scores based on responses to SHP2i and to SHP2i+MEKi. While KRAS(G12X) tumor cells were sensitive to the combination, RAS(WT) cells were also affected (**Fig. 3D**). These findings reinforce that the SHP2i+MEKi combination lacks tumor selectivity, likely explaining its poor tolerability in clinical trials and limiting its utility for treating KRAS(G12X) cancers.

### RAS(Q61R/K) cells are insensitive to combined SHP2i+MEKi due to feedback activation of RAS(Q61R/K) and induction of a SHP2 conformation with reduced SHP2i binding

We previously showed using a first generation SHP2i (SHP099) that tumor cells harboring RAS(Q61R/K) mutations exhibited complete resistance to SHP2i alone or in combination with MEKi^18^. While RAS(Q61R/K)-mutant tumors are generally considered SHP2-independent, their resistance to dual SHP2i+MEKi therapy was unexpected, as feedback RAS activation upon MEKi treatment is thought to occur through RAS(WT) and should therefore be sensitive to SHP2i. The question prompted us to further investigate the biochemical basis of RAS(Q61K/R)-driven resistance to combined SHP2i+MEKi.

First, we validated our previous result with TNO155^28^, a more potent, clinically relevant SHP2i, by treating SKMEL2 (NRASQ61R) cells with SHP2i, MEKi, or the combination. Consistent with our previous report^18^, MAPK rebound was fully resistant to combined SHP2i+MEKi (**Fig. 4A**). We next assessed feedback activation of RAS, by performing active RAS pulldown assays in SKMEL2 cells, which are homozygous for NRAS(Q61R). Following 24 hours of SHP2i+MEKi treatment, we observed elevated NRAS activity in response to the treatment (**Fig. 4B**), indicating that feedback activation of NRAS(Q61R), not RAS(WT), primarily drives adaptive MAPK recovery in these cells.

**Figure 4.**
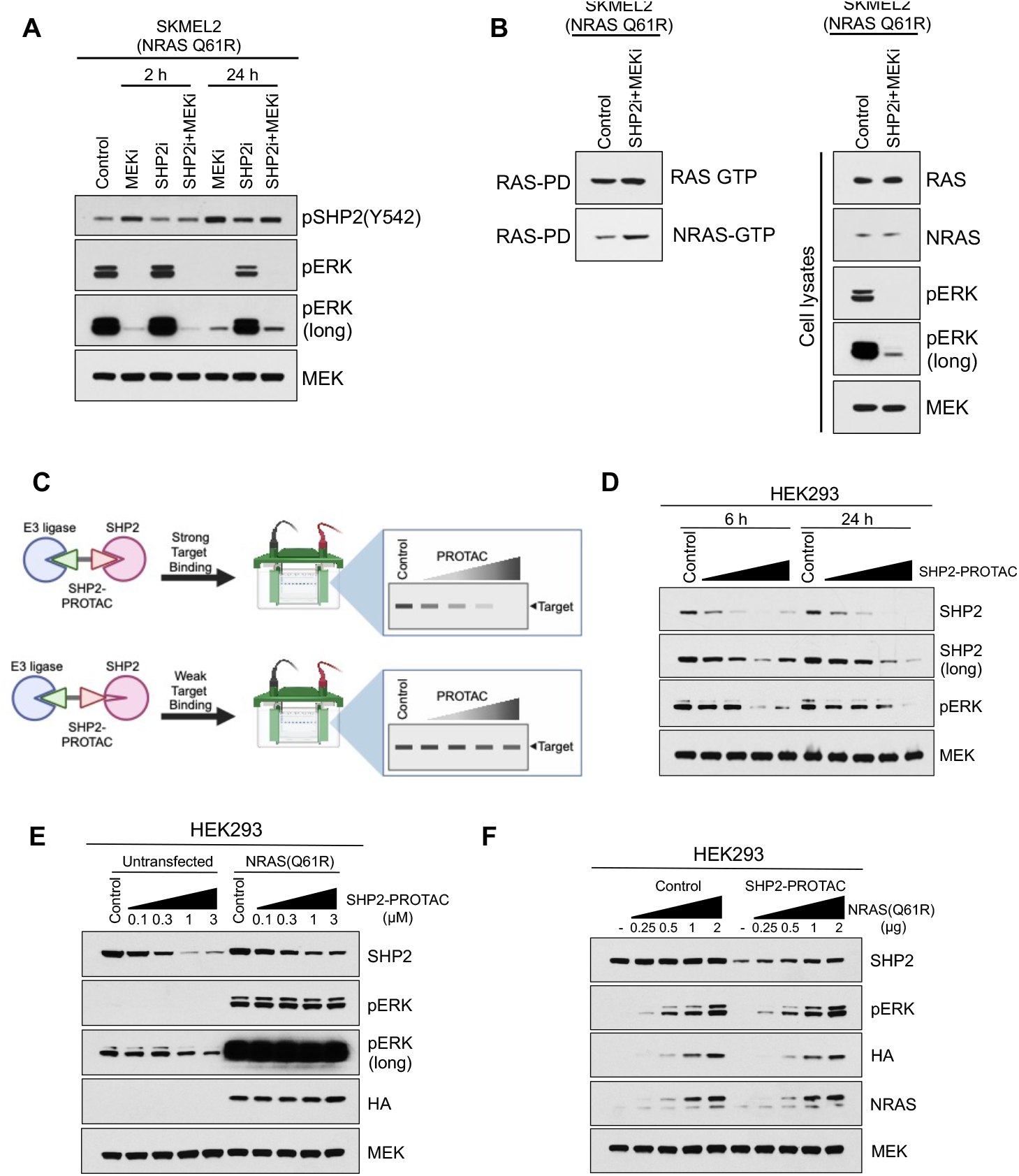
RAS(Q61R/K) cells are insensitive to combined SHP2i+MEKi treatment due to feedback activation of RAS(Q61R/K) and induction of a SHP2 conformation with reduced binding to SHP2i. **(A)** NRAS(Q61R) SKMEL2 cells were treated for 2 or 24 hours with either MEKi trametinib (5 nM), SHP2i TNO155 (5 µM), or the combination MEKi+SHP2i. SHP2 and MAPK pathway activation were monitored by immunoblotting, respectively indicated by pSHP2^Y542^ and pERK. **(B)** SKMEL2 cells were treated for 2 hours with SHP2i TNO155 (5 µM) combined with MEKi trametinib (5 nM). The level of activated RAS (RAS-GTP) and activated NRAS (NRAS-GTP) were assessed using a RAS-pull down assay. **(C)** Schematic representation of PROTAC-mediated targeted degradation of SHP2, as a proxy for the strength of binding of the warhead (SHP2i) to SHP2. Created in BioRender. Baars, B. (2025) https://BioRender.com/k62r551. (**D**) HEK293 cells were treated with increasing concentrations of the SHP2-PROTAC SHP2-D26 (0.1, 0.3, 1, 3 µM) for 6 or 24 hours. SHP2 degradation and MAPK pathway activation were assessed by immunoblotting. **(E)** HEK293 cells ectopically expressing HA-tagged NRAS(Q61R) were treated for 24 hours with increasing concentrations of the SHP2-PROTAC SHP2-D26. SHP2 degradation and MAPK pathway activation were assessed by immunoblotting. **(F)** HEK293 cells were transfected with the indicated amounts of HA-tagged NRAS(Q61R), followed by treatment with SHP2-PROTAC SHP2-D26 (0.5 µM) for 24 hours. SHP2 degradation and MAPK pathway activation were assessed by immunoblotting.

We next investigated whether NRAS(Q61R) activation affects binding of SHP2i by promoting a conformational change in SHP2. To test this, we used Proteolysis Targeted Chimera (PROTAC) mediated degradation (**Fig. 4C**) by a SHP2-PROTAC degrader^31^ as a proxy for the strength of SHP2i binding to SHP2. SHP2-PROTAC treatment significantly reduced SHP2 protein levels (**Fig. 4D**). However, when NRAS(Q61R) or KRAS(Q61K) were ectopically expressed, SHP2 degradation by SHP2 PROTAC was markedly reduced (**Fig. 4E, 4F, S3A, S3B**), indicating that RAS(Q61K/R) expression altered SHP2 conformation, impairing inhibitor accessibility. All of these findings support a mechanism of resistance to SHP2i+MEKi in RAS(Q61K/R) tumor cells, in which both feedback-driven RAS(Q61K/R) hyperactivation and conformational alterations in SHP2 contribute to evasion of SHP2-targeted therapies.

### Sensitivity to any single RAS-targeted inhibitor is restricted to an overlapping subset of RAS(MUT)-inhibitor sensitive tumors

In addition to panRAS-GEF(OFF)i (e.g., SHP2 and SOS1 inhibitors), panKRAS(OFF)i, as well as panRAS(ON)i have recently been developed. To assess their SII, we determined MAPK signaling and growth inhibition in a panel of cell line models, including normal cells and KRAS(G12C)- and KRAS(G12D) cancer lines with defined sensitivity to KRAS(G12C)- or KRAS(G12D)-specific inhibitors (**Fig. 5A, B**). We first compared the SIIs of a panRAS-GEF(OFF)i, a panKRAS(OFF)i, and a panRAS(ON)i, alongside the clinical MEKi trametinib. MEKi suppressed signaling and growth to varying degrees across these cell lines, without distinguishing between KRAS(G12X) and RAS(WT), indicating a low SII. These results are consistent with clinical data demonstrating modest efficacy and dose-limiting toxicities when MEKi is used in RAS(MUT) cancers^32-34^. Treatment with the panKRAS(OFF)i yielded a positive SII in KRAS(G12X) inhibitor-sensitive models but a substantially lower SII in cell lines insensitive to KRAS(G12X)-specific inhibitors. Treatment with the panRAS(ON)i resulted in MAPK and growth inhibition in both KRAS(MUT) and RAS(WT) cells, showing a positive, but low SII.

**Figure 5.**
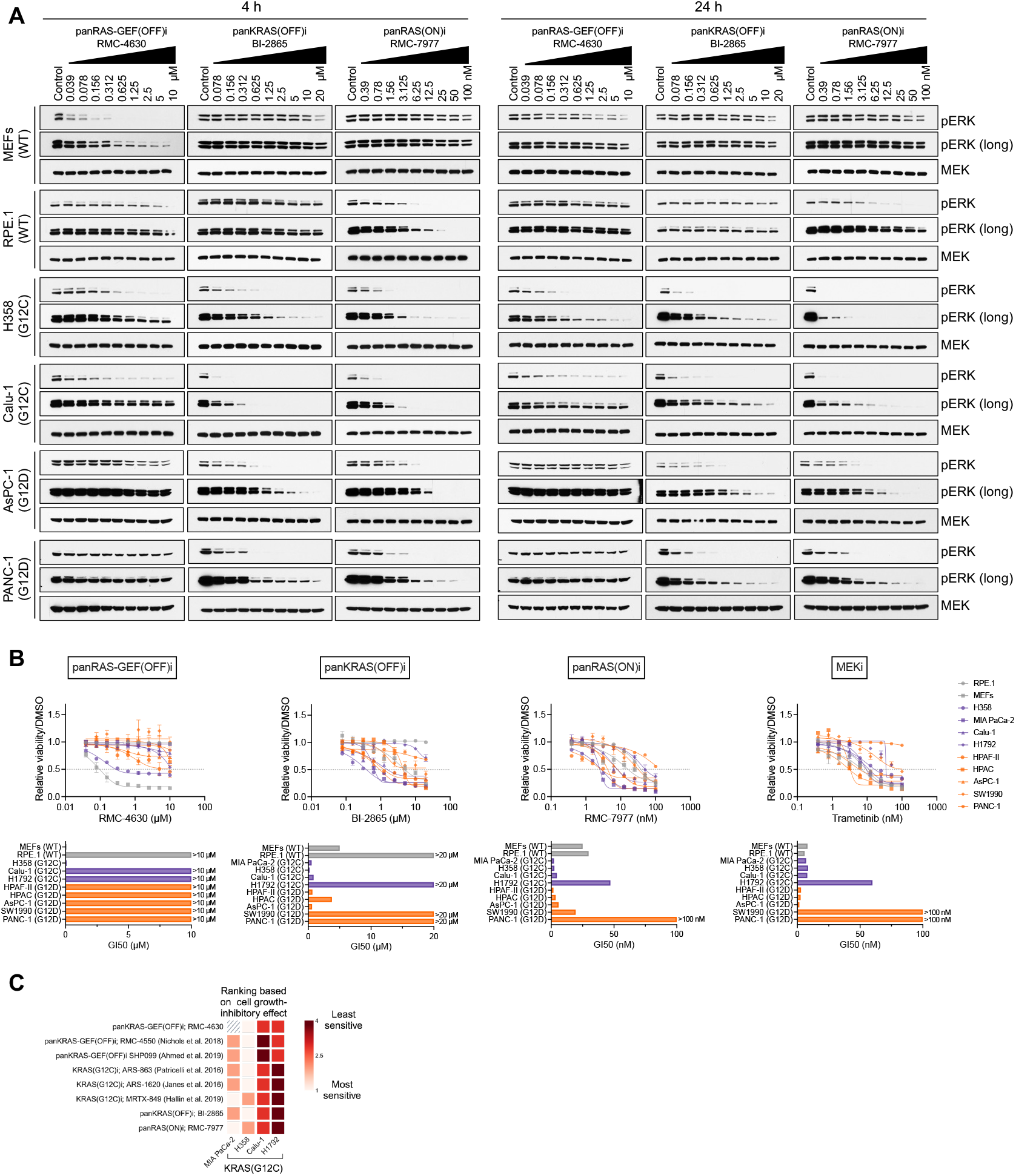
Sensitivity to any single RAS-targeted inhibitor is restricted to the subset of RAS(MUT)-specific inhibitor sensitive tumors. **(A)** The indicated RAS(WT), KRAS(G12C) and KRAS(G12D) cells were treated with increasing concentrations of panRAS-GEF(OFF)i RMC-4630, panKRAS(OFF)i BI-2865, or panRAS(ON)i RMC-7977. After 4 or 24 hours of treatment, MAPK pathway activation was assessed by immunoblotting, indicated by pERK. (**B**) Cell viability inhibition was assessed after 6 days of treatment. Dose-response curves were generated (top) and for each compound, GI50 for each cell line was generated and shown in the bar plot (bottom). (**C**) Comparative analysis of inhibitor sensitivity incorporating cell viability data from other studies. Four KRAS(G12C)-mutant cell lines that were used in our experiments and other studies were ranked based on their sensitivity to indicated inhibitors (most sensitive = 1, least sensitive = 4). Heat map was created in BioRender. Baars, B. (2025) https://BioRender.com/b56z167. Data are presented as mean ± SEM from three experimental replicates (n=3).

Finally, we carried out a comparative analysis of our cell growth inhibition data and previously published reports of KRAS(G12C) cell lines sensitivity to KRAS(G12C)i, panRAS-GEF(OFF)i, panKRAS(OFF)i, or panRAS(ON)i. We observed similar overall relative sensitivities across cell line models to different RAS inhibitors (**Fig. 5C**), supporting the notion that the high antitumor efficacy of RAS inhibitors is largely restricted to the same subset of RAS(MUT) cancers. Consequently, the majority of RAS(MUT) tumors are predicted to show limited responsiveness to any one paralog or state-selective RAS inhibitor.

## Discussion

RAS(MUT)-specific inhibitors have theoretically unlimited Therapeutic Index and thus can be clinically dosed at saturating concentrations. However, the Overall Response Rate (ORR) to these drugs is at best around 50%^9-11^. In addition to RAS(MUT)-specific inhibitors, several paralog and state-selective inhibitors targeting RAS activity have been developed as possible new therapeutic agents for RAS(MUT) cancers. These include indirect panRAS-GEF(OFF)i (e.g., SHP2i and SOS1i), direct panKRAS(OFF)i, sparing HRAS and NRAS (e.g., BI-2852, BI 3706674, ERAS-4001) and direct panRAS(ON)i (e.g., RMC-7977, RMC-6236, ERAS-0015, GFH547). In this study, we assessed MAPK pathway and growth inhibition by such RAS inhibitors in models of RAS(MUT) and RAS(WT) (normal) cells and compared their signaling inhibition index - SII (i.e. the differential inhibition of oncogenic signaling between tumor and normal cells) and relative growth inhibition. To represent normal RAS/MAPK signaling we used selected RAS(WT) models of varying sensitivity to MEKi and considered MAPK and growth inhibition by RAS signaling inhibitors in any of these models as suggesting potential on target/off tumor toxicity.

Consistent with previous reports, SHP2i showed activity in KRAS(G12X)-expressing cells. However, we found that KRAS(G12V/D) were less sensitive than KRAS(G12C/A/S). These results potentially explain the modest results of early phase clinical trials of SHP2i in patients with KRAS(G12X) tumors^35^.

We previously reported KRAS(G13D) cells to be resistant to SHP2i^18^, whereas others found them to be sensitive to SHP2i^15^ or SOS1i^20^. We found that differential expression of NF1 accounts for this discrepancy. KRAS(G13D/NF1-depleted) cells were completely insensitive to SHP2i or SOS1i, whereas KRAS(G13D/NF1-proficient) cells were modestly sensitive. These data are consistent with previous reports indicating KRAS(G13D) to be sensitive to NF1-mediated hydrolysis^36^ and therefore dependent on upstream RTK/SHP2/SOS signaling (EGFR) for its activity^37,38^.

Despite early preclinical findings suggesting that patients with RAS(G12X) tumors could benefit from SHP2i+MEKi therapy, clinical efficacy has been modest. We found that SHP2i+MEKi is highly potent not only in KRAS(G12X) but also in RAS(WT) cells, indicating a low SII, thus explaining the modest clinical results of the combination.

NRAS(Q61R) cells are resistant to SHP2i, however the basis of their resistance to combined SHP2i+MEKi^18^ was not obvious. NRAS(Q61R) is thought to be at the maximum of its activity and thus feedback MAPK activation should be carried through RAS(WT) activation, which is sensitive to SHP2i. However, we found that NRAS(Q61R) was hyperactivated in response to MAPK inhibition. This, together with SHP2 adopting a conformation with reduced SHP2i binding, as evidenced by decreased degradation by a SHP2 PROTAC, promoted resistance to SHP2i in the context of combined SHP2i+MEKi. As SHP2i bind the inactive, closed conformation of SHP2, RAS hyperactivation could promote a shift of the equilibrium of SHP2 towards the active conformation and thus reduce binding of SHP2i. Further studies are warranted to better characterize this conformational shift of SHP2, thus far only supported by cell-based studies.

By assessing in parallel SII and growth inhibition by panRAS-GEF(OFF)i, panKRAS(OFF)i and panRAS(ON)i in KRAS(G12X) and RAS(WT) (normal) cell models, we found that panKRAS(OFF)i show a positive SII, with minimal activity in normal cells. PanRAS(ON)i are active in both RAS(MUT) and RAS(WT) cells, with a positive, yet narrow SII. Importantly, all paralog- and state-selective RAS inhibitors are more potent in cell line models that are also sensitive to RAS(MUT)-specific inhibitors. This suggests that all RAS-targeting drugs to date are active in a similar “RAS inhibitor-sensitive” portion of tumors. These results emphasize the need for improved strategies to target the portion of RAS(MUT) patients that do not respond to any of the current RAS-targeting inhibitors.

Together our findings reinforce the notion that the effectiveness of paralog- and state-selective inhibitors depends largely on specific RAS mutations and cellular context (e.g. NF1 expression), and highlight the need to integrate SII considerations into the development and clinical application of RAS-targeted therapies.

## Author Contributions

B.B. designed and performed experiments, analyzed data and wrote the manuscript. A.O.R, B.G. X.Z., M.D. and C.A. performed experiments and analyzed data. S.A.A. designed experiments and analyzed data. S.W. developed and contributed the SHP2 PROTAC and analyzed data. E.G. designed experiments and analyzed data. P.I.P. conceived the study, supervised and coordinated research, analyzed data and wrote the manuscript.

## Acknowledgments

We thank Mallika Singh and Kyle Seamon for facilitating the provision of RMC-7977 and for valuable discussions. P.I.P. was supported by the US National Institutes of Health (**NIH**) (**R01 CA285713, R01 CA240362, R01CA238229**), a V Foundation Translational Grant, the Melanoma Research Alliance, the Breast Cancer Alliance, the Melanoma Research Foundation, the Irma T. Hirschl Trust and Tisch Cancer Institute developmental awards. E.G. was supported by **NIH** grants **R01CA223243** and **P30CA013330**.

## Material and Methods

### Compounds

TNO155 (S8987) and Trametinib (S2673) were obtained from Selleckchem. BI-3406 (HY-125817), MRTX-0902 (HY-145926) and RMC-4630 (HY-141523) were obtained from Medchem Express. BI-2865 (C-1443) was obtained from Chemgood. SHP2 degrader (SHP2-D26) was developed and contributed by Shaomeng Wang (University of Michigan). RM-042/RMC-7977 (HY-156498) was either contributed by Revolution Medicines or obtained from MedChem Express. Compounds were dissolved in DMSO to yield 10 mmol/L stock.

### Cell lines

H358, Calu-1, H1792, MIA PaCa-2, H1573, A549, SW620, HPAC, HPAF-II, AsPC-1, PANC-1, SW1990, HCT-15, HCT-116, DLD-1, SKMEL2, and HEK293T cell lines were purchased from American Type Culture Collection. HEK293 cell line was purchased from Life Technologies. Isogenic SW48 cells were obtained from Horizon Discovery. HeLa cells were kindly provided by Ramon Parsons (Icahn School of Medicine at Mount Sinai). RPE.1, AC16, HCT-E, and MRC5 cell lines were a gift from Anthony Faber (Virginia Commonwealth University). MEF cell lines has been described previously (7) All the cells used in the study were maintained in a humidified incubator at 37°C with 5% CO2 and cultured in RPMI 1640 (ThermoFisher, 11-875-119) or DMEM (ThermoFisher, 10313-039) supplemented with 10% FBS (Biowest, S1680), 2 mmol/L glutamine (ThermoFisher, 25-030-164), and 100 IU/mL penicillin/streptomycin (ThermoFisher, 15-140-163) and were passaged from three to five times. Cell lines regularly tested negatively for Mycoplasma using the Venor GeM Mycoplasma Detection Kit (Sigma).

### Plasmids and transfections

pMEVHA-KRAS2A-WT was purchased from Biomyx (P1020). NRAS was subcloned using pMEV as a backbone. N-terminal HA-tagged KRAS was subcloned into pTRI-BlasticidinS plasmid (From Yuki Otsuka). KRAS (G12C, G12D, G12A, G12S, G12V, Q61K) and NRAS (Q61R) activating mutants were generated with the QuikChange II XL Site-Directed Mutagenesis Kit (Agilent, 200522). Transfections were carried out using Lipofectamine 2000 (ThermoFisher, 11668019).

### NF1 gene silencing

DLD-1 cells and HeLa cells were transfected with 600 pmol of ON-TARGETplus human NF1 siRNA (Horizon Discovery) or ON-TARGETplus non-targeting control pool (Horizon Discovery) using Lipofectamine RNAiMAX transfection reagent (Invitrogen) following manufacturer’s instructions. 24 hr later, cells were treated with the indicated drugs, and NF1 levels were evaluated by Western blot as previously described.

### Lentivirus transduction and doxycycline-inducible stable cell line generation

Lentivirus was produced by transfecting HEK293T cells with the pTRI-BlaS transfer plasmid containing various KRAS variants (KRAS (WT, G12C, G12D, G12S, G12A, G12V, Q61K), the packaging vector psPAX2 (Addgene), and envelope vector pMD2.G (Addgene) a 5:4:1 ratio using Lipofectamine 2000 (ThermoFisher, 11668019). The viral supernatant was collected 72 hours after transfection and filtered through a 0.45-μm filter unit (MilliporeSigma, SLHPR33RS). After 24 h, HEK293T medium was replaced with fresh medium, and 48 h after the co-transfection, the medium containing lentiviral particles was collected, filtered, and used to infect HeLa cells after the addition of 10 μg/mL polybrene (Sigma-Aldrich). HeLa cells were incubated overnight at 37°C prior to the removal of viral particles and the addition of fresh media. Transduced cells were then selected for by culturing in the presence of 3 µg/ml Blasticidin (Invivogen, ant-bl-05) for approximately 10 days. Cells were cultured in the presence of 50-600 ng/ml doxycycline for 24 hours prior to use in experiments.

### Western blot and active RAS pull-down

Cells were washed with PBS and lysed on ice for 5 minutes in NP40 buffer (50 mmol/L Tris pH 7.5, 1% NP40, 150 mmol/L NaCl, 10% Glycerol 1 mmol/L EDTA) supplemented with protease (Roche, 11873580001) and phosphatase inhibitors (Roche, 4906837001). Lysates were centrifuged at 15,000 rpm for 10 minutes, and the protein concentration was quantified using BCA (ThermoFisher, 23221, 23224). Proteins were separated by NuPAGE and 4% to 12% Bis Tris Gel (Invitrogen, NP0321BOX), and they were immunoblotted and transferred to nitrocellulose membranes (GE Healthcare, 45-004-006) according to standard protocols. Membranes were immunoblotted overnight with antibodies against: pMEK1/2^Ser217/22^ (9154), MEK1 (2352), pERK1/2^Thr202/Tyr204^ (4370) from Cell Signaling; SHP2 (sc-7384) and NRAS (sc-31) from Santa Cruz Biotechnology; pSHP2^Y542^ from abcam (ab62322); NF1 from Invitrogen (PA5-103218), hemagglutin (HA) from Biolegend (901501); V5 from ThermoFisher (R960-25); KRAS from (Proteintech, 12063-1-AP). The next day, membranes were probed with anti-rabbit IgG (Cell Signaling, 7074) or anti-mouse IgG secondary antibody (Cell Signaling, 7076), and chemiluminescent signals were detected on X-ray films.

RAS pull-down kit (ThermoFisher, 16117) was used to determine levels of RAS-GTP, according to the manufacturer’s protocol. RAS was detected by western blot using either an antibody for KRAS (Santa Cruz Biotechnology, sc-30), or pan-RAS antibody provided with the kit.

### Cellular viability assay

Cells were plated at a density of 0.5-1 × 10^3^ cells per well in 96-well plates (Corning™, 3595). The next day, cells were treated with inhibitors as indicated in regular growth media for 6 days. 10 µl of Cell Counting Kit-8 (Dojindo Molecular Technologies, Inc., CK04) was added, and the cells were incubated at 37 °C for 2.5 hours. Cell viability was then determined by measuring the absorbance at 450 nm using a plate reader. Dose response curves and GI50 values were calculated using log-transformed, normalized data in Graphpad Prism 10.4.1.

## Figure legends

**Supplementary Figure 1.**
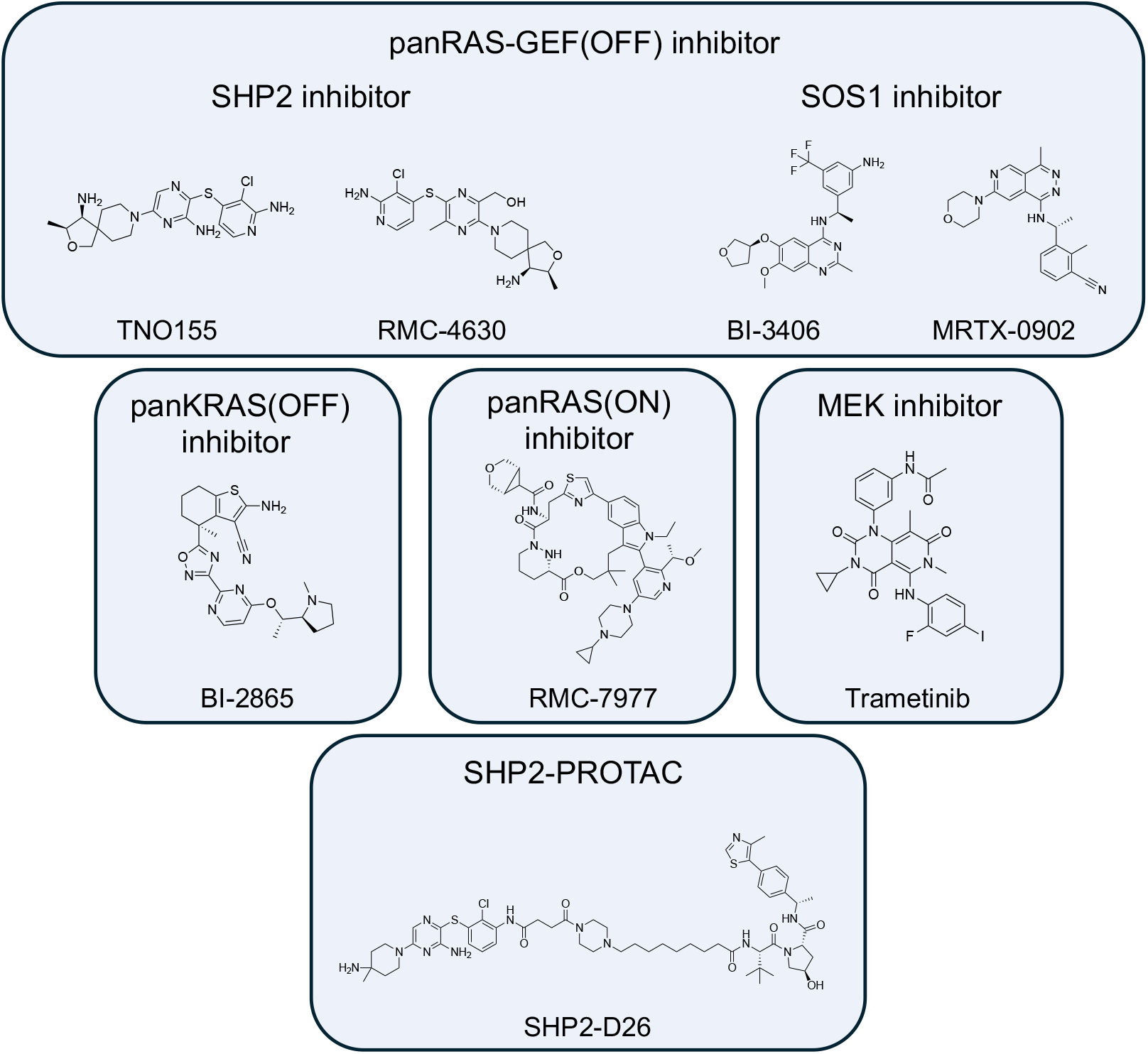
Chemical structures of compounds used in this study.

**Supplementary Figure 2.**
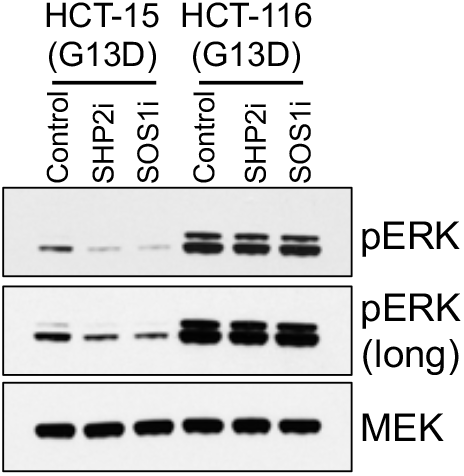
KRAS(G13D) cells are insensitive to next generation SHP2i and SOS1i. KRAS (G13D)-mutant cell lines HCT-15 and HCT-116 were treated for 2 hours with SHP2i TNO155 (5 µM) or SOS1i BI-3406 (10 µM). MAPK pathway activity was assessed by immunoblotting.

**Supplementary Figure 3.**
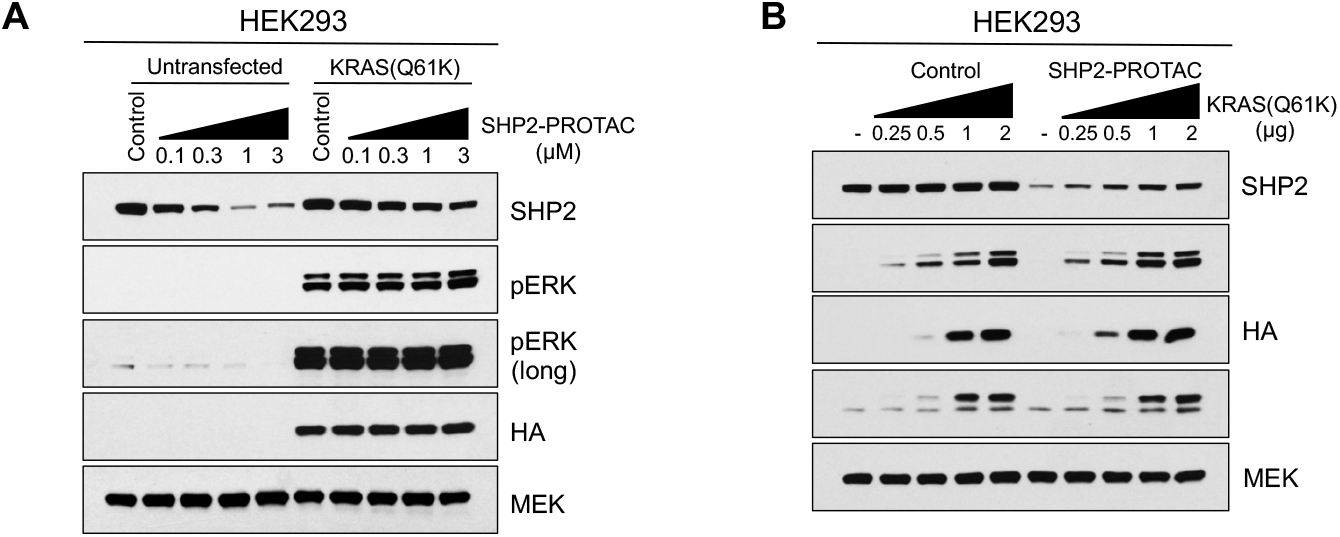
KRAS(Q61K)-expressing cells induce altered SHP2 conformation unable to bind SHP2 PROTAC. **(A)** HEK293 cells ectopically expressing HA-tagged KRAS (Q61K) were treated for 24 hours with increasing concentrations of SHP2-PROTAC SHP2-D26. SHP2 degradation and MAPK pathway activation were assessed by immunoblotting. (**B**) HEK293 cells were transfected with indicated amounts of HA-tagged KRAS (Q61K), followed by treatment with SHP2-PROTAC SHP2-D26 (0.5 µM) for 24 hours. SHP2 degradation and MAPK pathway activation were assessed by immunoblotting.

